# *In vivo* microscopy reveals the impact of *Pseudomonas aeruginosa* social interactions on host colonization

**DOI:** 10.1101/488262

**Authors:** Chiara Rezzoagli, Elisa T. Granato, Rolf Kümmerli

## Abstract

Pathogenic bacteria engage in social interactions to colonize hosts, which include quorum-sensing-mediated communication and the secretion of virulence factors that can be shared as “public goods” between individuals. While *in-vitro* studies demonstrated that cooperative individuals can be displaced by “cheating” mutants freeriding on social acts, we know less about social interactions in infections. Here, we developed a live imaging system to track virulence factor expression and social strain interactions in the human pathogen *Pseudomonas aeruginosa* colonizing the gut of *Caenorhabditis elegans*. We found that shareable siderophores and quorum-sensing systems are expressed during infections, affect host gut colonization, and benefit non-producers. However, non-producers were unable to cheat and outcompete producers. Our results indicate that the limited success of cheats is due to a combination of the down-regulation of virulence factors over the course of the infection, the fact that each virulence factor examined contributed to but was not essential for host colonization, and the potential for negative-frequency dependent selection. Our findings shed new light on bacterial social interactions in infections and reveal potential limits of therapeutic approaches that aim to capitalize on social dynamics between strains for infection control.

## Introduction

During infections, pathogenic bacteria secrete a wide range of extracellular virulence factors to colonize and grow inside the host [1, 2]. Secreted molecules include siderophores for iron scavenging, signaling molecules for quorum sensing (QS), toxins to attack host cells, and matrix compounds for biofilm formation [3–6]. *In-vitro* studies have shown that extracellular virulence factors can be shared as “public goods” between cells, and thereby benefit individuals other than the producing cell [7–9]. There has been enormous interest in understanding how this form of bacterial cooperation can be evolutionarily stable since secreted public goods can be exploited by non-cooperative mutants called “cheats”, which do not engage in cooperation yet still benefit from the molecules produced by others [10–12].

There is increasing evidence that social interactions and cooperator-cheat dynamics might also matter within hosts [9, 13, 14]. For instance, in controlled infection experiments engineered non-producers, deficient for the production of specific virulence factors, could outcompete producers and thereby reduce virulence [15–19], but in other cases the success of non-producers was compromised [20, 21]. Other studies have followed chronic human infections within patients over time and reported that virulence-factor-negative mutants emerge and spread, with the mutational patterns suggesting cooperator-cheat dynamics [7, 22, 23]. These findings spurred ideas of how social interactions within hosts could be manipulated for therapeutic purposes [14, 24, 25]. Suggested approaches include: inducing cooperator-cheat dynamics to steer infections towards lower virulence [7, 26]; introducing less virulent strains with medically beneficial alleles into established infections [24]; and targeting secreted virulence factors to control infections and constrain the evolution of resistance [25, 27–31].

However, all these approaches explicitly rely on the assumption that the social traits of interest are: (i) expressed inside hosts; (ii) important for host colonization; (iii) exploitable; and (iv) induce cooperator-cheat dynamics as observed *in vitro* [9] – assumptions that have not yet been tested in real time inside living hosts. Here, we explicitly test the importance of bacterial social interactions within hosts by using *in-vivo* fluorescence microscopy to monitor bacterial virulence factor production, host colonization and strain interactions, using the opportunistic pathogen *Pseudomonas aeruginosa* and its nematode model host *Caenorhabditis elegans* [32–34].

*C. elegans* naturally preys on bacteria [35]. While most bacteria are killed during ingestion, a small fraction of cells survives [36], which can, in the case of pathogenic bacteria, establish an infection in the gut [37]. *P. aeruginosa* deploys an arsenal of virulence factors that facilitate successful host colonization [38]. For example, the two siderophores pyoverdine and pyochelin scavenge host-bound iron during acute infections to enable pathogen growth [6, 39–41]. *P. aeruginosa* further secretes the protease elastase, the toxin pyocyanin, and rhamnolipid biosurfactants to attack host tissue [7, 42, 43]. Production of these latter virulence factors only occurs at high cell densities and is controlled by the Las and the Rhl QS-systems [44]. Because both QS-regulated virulence factors and siderophores were shown to be involved in *C. elegans* killing [37, 45–48], we used them as focal traits for our study.

To tackle our questions, we first conducted experiments with fluorescently tagged *P. aeruginosa* bacteria (PAO1) to follow infection dynamics, from the first uptake through feeding up to a progressed state of gut infection. We then constructed promoter-reporter fusions for genes involved in the synthesis of the two siderophores (pyoverdine and pyochelin) and the two QS-regulators (LasR and RhlR) to track *in vivo* virulence factor gene expression during host colonization. Subsequently, we used mutant strains deficient for virulence factors to determine whether they show compromised colonization abilities. Finally, we followed mixed infections of wildtype and mutants over time to determine the extent of strain co-localization in the gut, and to test whether secreted virulence factors are indeed exploitable by non-producers in the host.

## Material and methods

### Strain and bacterial growth conditions

Bacterial strains, primers and plasmids used in this study are listed in the Supplementary Tables S1-S3. Details on strain construction are given in the Supplementary Methods. For all experiments, overnight cultures were grown in 8 ml Lysogeny broth (LB) in 50 ml tubes, incubated at 37°C, 220 rpm for 18 hours, washed with 0.8% NaCl solution and adjusted them to OD_600_ = 1. Nematode Growth Media (NGM) plates [0.25% Peptone, 50 mM NaCl, 25mM [PO_4_^-^], 5 μg/ml Cholesterol, 1mM CaCl_2_, 1mM MgSO_4_ supplemented with 1.5% agar, 6 cm diameter] were seeded with 50 μl of bacterial culture and incubated at 25°C for 24 hours. All *P. aeruginosa* strains used in this study showed equal growth on NGM exposure plates (Supplementary Figure S1). Peptone was purchased from BD Biosciences, Switzerland, all other chemicals from Sigma Aldrich, Switzerland.

### Nematode culture

We used the temperature-sensitive, reproductively sterile *C. elegans* strain JK509 (*glp-1*(*q231*) III). This strain reproduces at 16°C, but does not develop gonads and is therefore sterile at 25°C. Worms were maintained fertile at 16°C on High Growth Media (HGM) agar plates (2% Peptone, 50 mM NaCl, 25mM [PO_4_^-^], 20 μg/ml Cholesterol, 1mM CaCl_2_, 1mM MgSO_4_) seeded with the standard food source *E. coli* strain OP50 [49]. For age synchronization, plates were washed with sterile distilled water and adult worms were killed with hypochlorite-sodium hydroxide solution to isolate eggs [49]. Eggs were placed in M9 buffer (20 mM KH_2_PO_4_, 40 mM Na_2_HPO_4_, 80 mM NaCl, 1 mM MgSO_4_) and incubated at 16°C for 16-18 hours to hatch. Then, L1 larvae were transferred to OP50-seeded HGM plates and incubated at 25°C for 28 hours to reach L4 developmental stage. Worms and OP50 bacteria were provided by the Caenorhabditis Genetic Center (CGC), which is supported by the National Institutes of Health - Office of Research Infrastructure Programs (P40 OD010440).

### *C. elegans* infection protocol

Synchronized L4 worms were washed from HGM plates with M9 buffer + 50 μg/ml kanamycin (M9-Kan), and washed three times with M9-Kan for worm surface-disinfection. Viable worms were further separated from any debris by sucrose flotation [50] and rinsed three time in M9 buffer to remove sucrose. The worm handling protocol for the main experiments is depicted in Figure 1A. Specifically, approximately 200 worms were moved to seeded NGM plates and incubated for 24 hours at 25°C. After this period of exposure to pathogens, infected worms were extensively washed with M9 buffer + 50 μg/ml chloramphenicol (M9-Cm) followed by M9 buffer, subsequently transferred to individual wells of a 6-well plate filled with sterile M9 buffer + 5 μg/ml cholesterol (M9+Ch Buffer) where they were kept up to 48 hours post exposure (hpe) and imaged after 0, 6 and 30 hpe.

**Figure 1.**
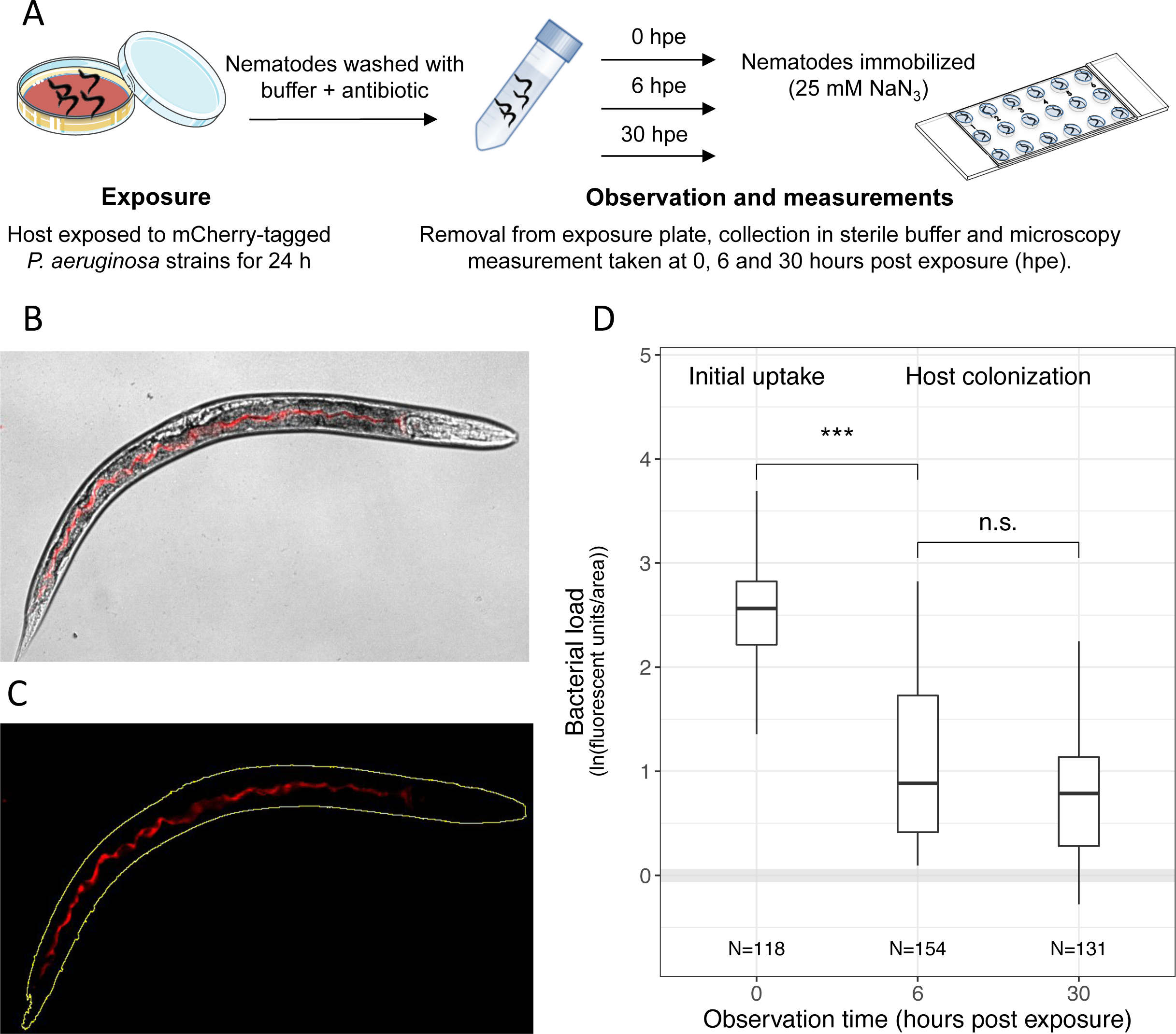
Quantifying *P. aeruginosa* infections in the *C. elegans* gut. (A) Experimental procedure: we used fluorescently tagged *P. aeruginosa* strains to examine bacterial colonization of the *C. elegans* gut. Per experiment, we exposed approximately 200 *C. elegans* nematodes to a lawn of mCherry-tagged PAO1 strains for 24 hours. Subsequently, nematodes were removed from the bacterial plate, surface washed and collected in sterile buffer for monitoring. After 0, 6, or 30 hours post exposure (hpe), approximately 30 nematodes were immobilized and transferred to microscopy slides for imaging. (B) Brightfield and fluorescence channel merged image depicting mCherry-fluorescent bacteria inside the host gut. (C) Bacterial load inside the nematode was quantified as the sum of fluorescence intensity across pixels in the region <u>o</u>f interest “ROI” (yellow outline) and standardised by total worm area. (D) Colonization dynamics of the wildtype strain PAO1-*mCherry*: immediately after removal from the exposure plate (0 hpe), worms showed high bacterial loads inside their guts. Bacterial load first declined when the worms where kept in buffer for 6 hours, but then remained constant for the next 24 hours. Grey shaded area indicates background fluorescence (mean +/- standard deviation) of worms exposed to the non-fluorescent, non-pathogenic *E.coli* OP50. N = number of worms from four independent experiments. *** = p < 0.001, n.s. = not statistically significant.

### Nematode survival assay

Our goal was to observe infections inside living hosts. To verify that worms stayed alive during the experiment (up to 48 hpe), we tracked the survival of infected population (50-90 worms) in M9-Ch buffer. Worms were observed for motility at 0, 24 and 48 hpe, by prodding them with a platinum wire. Worms were considered dead when they no longer responded to touches. Each bacterial strain was tested in three replicates and three independent experiments were carried out. We used *E. coli* OP50 as a negative control for killing. During this observation period, worms experienced only negligible killing by the colonizing bacteria, and we found no significant difference in killing between the non-pathogenic *E. coli* food strain and the *P. aeruginosa* strains (Supplementary Figure S2).

### Microscopy setup and imaging

For microscopy, we picked individual worms from the M9+Ch buffer and paralyzed them with 25 mM sodium azide before transferring them to an 18- well μ-slide (Ibidi, Germany). All experiments were carried out at the Center for Microscope and Image Analysis of the University Zürich (ZMB). For the colonization experiment, images were acquired on a Leica LX inverted widefield light microscope system with Leica-TX2 filter cube for mCherry (emission: 560 ± 40 nm, excitation: 645 ± 75 nm, DM = 595) and Leica-DFC-350-FX, cooled fluorescence monochrome camera (resolution: 1392 × 1040 pixels) for image recording (16-bit color depth). For gene expression experiments, microscopy was performed on the InCell Analyzer 2500HS (GE Healthcare) automated imaging system, using polychroic beam splitter BGRFR_2 (for mCherry, excitation: 575 ± 25 nm, emission: 607.5 ± 19 nm) and PCO – sCMOS camera (resolution: 2048 x 2048 pixels, 16-bit).

### Image processing and analysis

To extract fluorescence measurements from individual worms, images were segmented into objects and background, using an automated image segmentation workflow with *ilastik* software [51]. Segmented images were then imported in *Fiji* [52] to determine the fluorescence intensity (as “Raw Integrated Density”, i.e. the sum of pixels values in the selection) and area of each worm. Images obtained from the InCell microscope entailed 64 frames (8×8 grid) with 10% overlap. These frames were stitched together using a macro-automated version of the Stitching plugin in *Fiji* [53] prior to segmentation and analysis. To correct for background and host-tissue autofluorescence, we imaged, at each time point, worms infected with non-fluorescent strains (i.e. OP50 or PAO1), and used the mean intensity of these control infections to subtract background fluorescence values from worms infected with fluorescent strains.

### Competition assay in the host

For *in-vivo* competitions between PAO1-*mCherry* and PAO1Δ*pvdD*Δ*pchEF* or PAO1ΔlasR, overnight monocultures were washed twice with 0.8% NaCl solution, adjusted to OD_600_ = 1 and mixed at a 1:1 ratio. To control for fitness effects of the mCherry marker, we also competed PAO1-*mCherry* against the untagged PAO1. NGM plates were then seeded with 50 μl of mixed culture and incubated at 25°C for 24 hours. Worms were exposed to the mix for 24 hours and then recovered as previously described. After 6 and 48 hours post-exposure, individual worms were picked, immobilized with sodium azide and washed for 5 minutes with M9 + 0.003% NaOCl. Worms were washed twice with M9 buffer. We then transferred each individual worm to a 1.5 ml screw-cap micro tube (Sarstedt, Switzerland) containing sterilized glass beads (1 mm diameter, Sigma Aldrich). Worms were disrupted using a bead-beater (TissueLyser II, QIAGEN, Germany), shaking at 30 Hz for 1.5 min before flipping the tubes and shaking for an additional 1.5 min to ensure even disruption (adapted from [54]). Tubes were then centrifuged at 2000 x g for 2 min, the content was re-suspended in 200 μl of 0.8% NaCl and plated on two LB 1.2 % agar plates for each sample. Plates were incubated overnight at 37°C and left at room temperature for another 24 h to allow the fluorescent marker to fully mature. We then distinguished between fluorescent and non-fluorescent colonies using a custom built fluorescence imaging device (*Infinity 3* camera, Lumenera, Canada). We then calculated the relative fitness of PAO1-*mCherry* as ln(v)=ln{[*a*_48_×(1−*a*_6_)]/[*a*_6_×(1−*a*_48_)]}, where *a*_6_ and *a*_48_ are the frequency of PAO1-*mCherry* at 6 and 48 hours after recovery, respectively [55]. Values of ln(v)<0 or ln(v)>0 indicate whether the frequency of PAO1-*mCherry* increased (ln(v)< 0) or decreased (ln(v)>0) relative to its competitor.

### Co-localization analysis

To determine the degree of co-localization of two different bacterial strains in the host, we transferred nematodes to NGM plates seeded with a 1:1 mix of PAO1*-gfp* with either PAO1-*mCherry*, PAO1*ΔpvdDΔpchEF–mCherry*, or *PAO1ΔpvdDΔpchEF–mCherry*, or PAO1*ΔlasR-mCherry*. After a 24 hours grazing period, we picked single worms and imaged both mCherry- and GFP channels, using the InCell Analyzer 2500HS microscope. We used *Fiji* to straighten each worm with the *Straighten* plugin [56], and extracted fluorescence intensity values in the GFP and mCherry channels for each pixel from tail (X = 0) to head (X = 1) of the worm. To ensure that we only measure areas where bacteria were present, we restricted our analysis to the region of the worm gut, where bacterial colonization takes place. We then calculated Spearman correlation coefficients between the fluorescent signals, as a proxy for strain co-localization using RStudio v. 3.3.0 [57].

### Statistical analysis

All statistical analyses were performed with RStudio. We used Pearson correlations to test for associations between PAO1-*mCherry* fluorescence intensities and (a) recovered bacteria from the gut; and (b) total bacterial load in mixed infections. We used analysis of variance (ANOVA) to compare fluorescence values between observation times, strains and for comparisons to non-fluorescent controls. *P*-values were corrected for multiple comparisons using the post-hoc Tukey HSD test. To compare promoter expression data between PAO1 WT and mutant strains, and to compare relative fitness values between competitors in the competition assay, we used Welch’s two-sample t-test. To measure co-localization, we calculated the Spearman correlation coefficient ***ρ*** between the intensity of mCherry and GFP signals across the worm gut, and used ANOVA to test for differences between treatments.

## Results

### PAO1 colonization dynamics in the *C. elegans* gut

For all infection experiments, we followed the protocol depicted in Figure 1A-C. We first exposed worms to *P. aeruginosa* for 24 hours on NGM plates. Subsequently, worms were removed, washed, and treated with antibiotics to kill external bacteria. We then imaged infected worms under the microscope at different time points and quantified bacterial density and gene expression using fluorescent mCherry markers. We first confirmed that mCherry fluorescence is a suitable proxy for the number of live bacteria in *C. elegans*, by comparing fluorescence intensities in whole worms (Figure 1B) to the number of live bacteria recovered from the worms’ gut. Fluorescence intensity values positively correlated with the bacterial load inside the nematodes, both immediately after recovering the worms from the exposure plates and at 6 hours post exposure (hpe; Supplementary Figure S3, Pearson correlation coefficient at 0 hpe: r = 0.49, t_28_ = 3.02, p = 0.0053; at 6 hpe: r = 0.713, t_23_ = 4.88, p < 0.0001). As our goal was to image infections in living hosts, we further confirmed that worms stayed alive during the observation period (Supplementary Figure S2).

When following host colonization by PAO1-*mCherry* over time, we observed that immediately after removal from the exposure plate, worms carried large amounts of bacteria in their gut (Figure 1D). Subsequently, bacterial load significantly declined when worms were kept in buffer for 6 hours (ANOVA: t_391_ = −8.55, p < 0.001) and then remained constant for the next 24 hours (t_391_ = 0.61, p = 0.529). This pattern suggests that a large number of bacteria are taken up during the feeding phase, followed by the shedding of a high proportion of cells, leaving behind a fraction of live bacteria that establishes an infection and colonizes the worm gut.

### PAO1 expresses siderophore biosynthesis genes and QS regulators in the host

We then examined whether genes involved in the synthesis of pyoverdine (*pvdA*) and pyochelin (*pchEF*), and the genes encoding the QS-regulators *lasR* and *rhlR*, are expressed inside hosts. Worms were exposed to four different PAO1 strains, each carrying a specific promoter-*mCherry* fusion. Imaging after the initial uptake phase (0 hpe) revealed that, with the exception of *pchEF*, all genes were significantly expressed in the host (Figure 2; ANOVA, comparisons to the non-fluorescent control, for *pvdA*: t_754_ = 4.23, p < 0.001; for *pchEF*: t_754_ = 0.74, p = 0.461; for *lasR*: t_754_ = 2.96, p = 0.003; for *rhlR*: t_754_ = 10.37, p <0.001). Although fluorescence intensity declined over time (linear model, F_1,1795_ = 48.98, p < 0.001), we observed that apart from *pchEF*, all genes were still significantly expressed during the subsequent colonisation of the host at 30 hpe. (Figure 2; ANOVA, for *pvdA*: t_754_ = 4.87, p < 0.001; for *pchEF*: t_754_ = 0.684, p = 0.461; for *lasR*: t_754_ = 3.01, p = 0.003; for *rhlR*: t_754_ = 16.68, p < 0.001).

**Figure 2.**
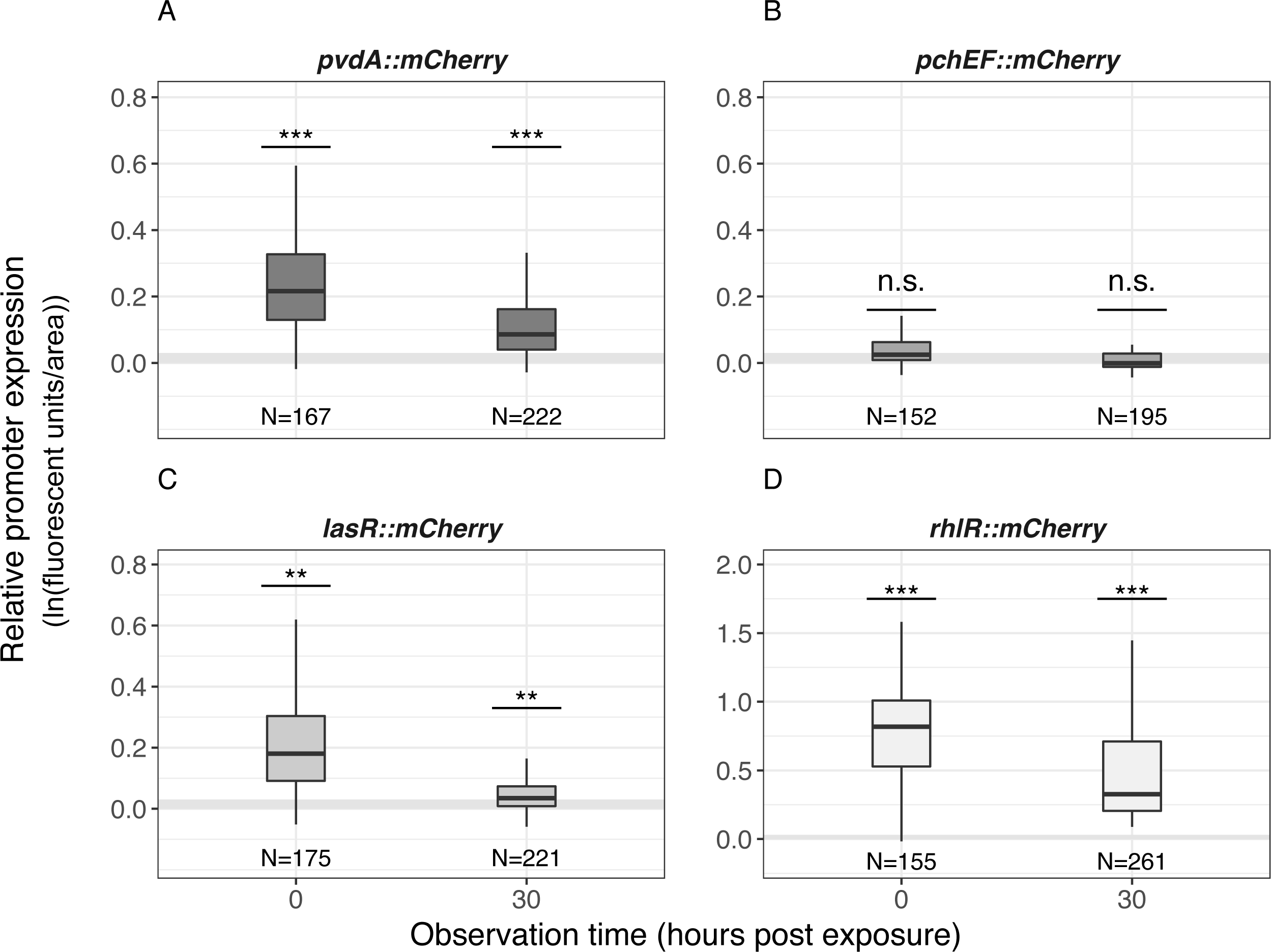
*P. aeruginosa* expresses genes for pyoverdine synthesis and quorum sensing regulators in the host gut. To quantify the expression of virulence factor genes inside hosts, worms were exposed to four PAO1 strains, each containing a promoter::*mCherry* fusion for either *pvdA (*pyoverdine synthesis*), pchEF* (pyochelin synthesis), *lasR* or *rhlR* (quorum sensing regulators). With the exception of *pchEF*, all genes were significantly expressed in the host, both at 0 and 30 hpe. Expression levels were standardised for bacterial load. Grey shaded areas depict background fluorescence (mean +/- standard deviation) of worms exposed to the non-fluorescent, non-pathogenic *E.coli* OP50. N = number of worms from four independent experiments. * = p < 0.05; ** = p < 0.01, *** = p < 0.001, n.s. = not statistically significant.

### Regulatory links between social traits operate inside the host

We know that regulatory links exist between the virulence traits studied here. While pyoverdine synthesis suppresses pyochelin production under stringent iron limitation [58], the Las-QS system positively activates the Rhl-QS system [44]. To test whether these links operate inside the host, we measured gene expression of each trait in the negative background of the co-regulated trait (Figure 3). For *pvdA*, we observed significant gene expression levels in both the wildtype PAO1 and the pyochelin-deficient PAO1Δ*pchEF* strain (Figure 3A), albeit the overall expression was slightly reduced in PAO1Δ*pchEF* (t-test, t_253_ = 8.67, p < 0.001). For *pchEF*, expression patterns confirm the suppressive nature of pyoverdine: the pyochelin synthesis gene was not expressed in the wildtype but significantly upregulated in the pyoverdine-deficient PAO1Δ*pvdD* strain (Figure 3B; t_296_ = −19.68, p < 0.001). For *lasR*, we found that gene expression was not significantly different in wildtype PAO1 compared to the Rhl-negative mutant PAO1Δ*rhlR*, confirming that the Las-QS system is at the top of the hierarchy and not influenced by the Rhl-system (Figure 3C; t_211_ = −1.50, p = 0.136). Conversely, the expression of *rhlR* was strongly dependent on a functional Las-system, and therefore only expressed in PAO1, but repressed in the Las-negative mutant PAO1Δ*lasR* (Figure 3D; t_156_ = 19.04, p < 0.001). These results show that (i) iron-limitation is strong in *C. elegans* as PAO1 primarily invests in the more potent siderophore pyoverdine; (ii) pyochelin can have compensatory effects when pyoverdine is lacking; and (iii) the loss of the Las-system leads to the concomitant collapse of the Rhl-system.

**Figure 3.**
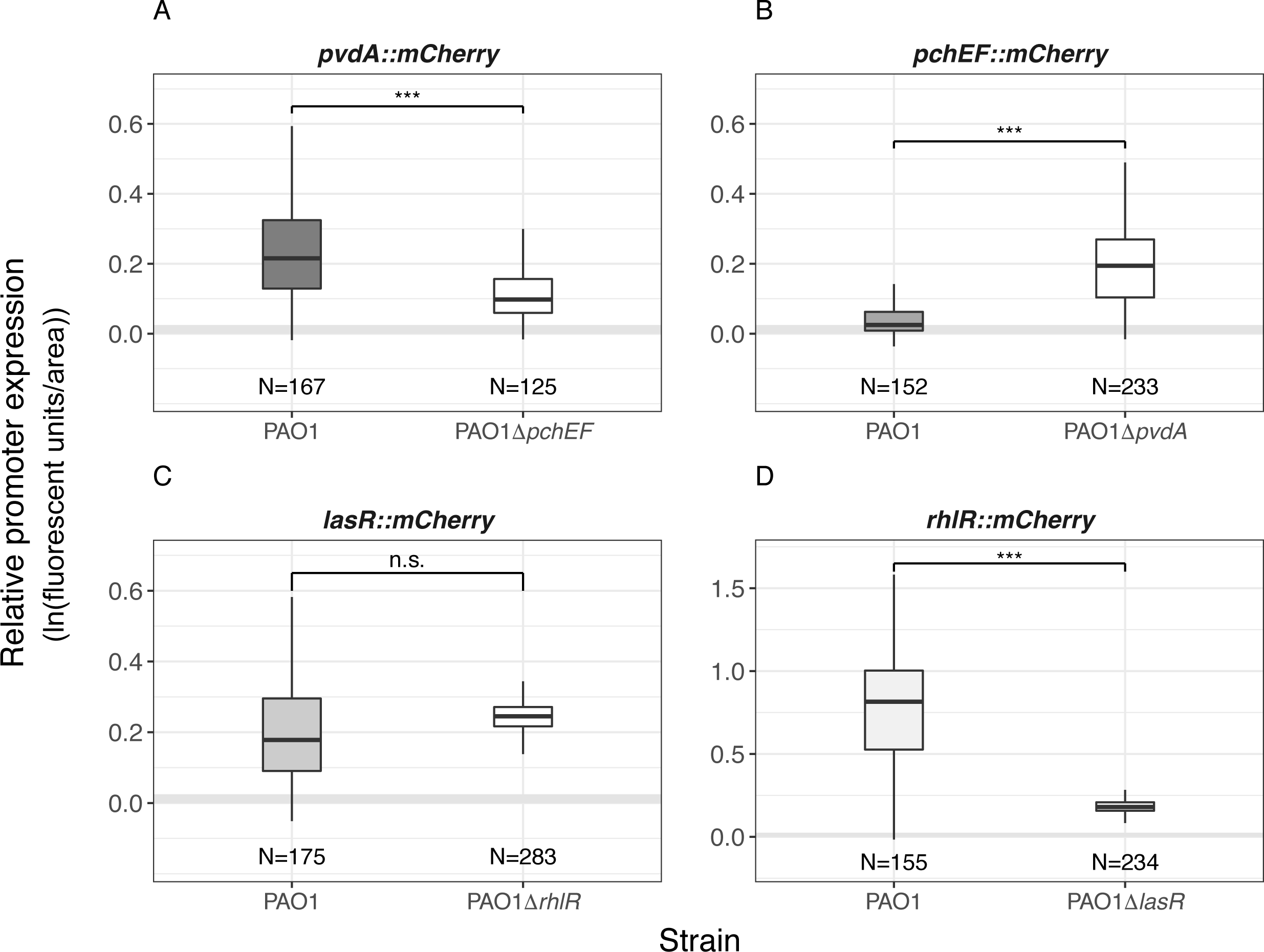
*P. aeruginosa* can switch between siderophores, while quorum sensing regulators act hierarchically. Because virulence traits are linked at the regulatory level, we measured gene expression of each trait in the negative background of the co-regulated trait. (A) The expression of the pyoverdine synthetic gene *pvdA* is significantly expressed in the wildtype and the pyochelin-negative background, but slightly reduced in the latter. (B) The pyochelin synthetic gene *pchEF* is significantly expressed in the pyoverdine-negative background, but silent in the wildtype. (C) The expression of the QS-regulator gene *lasR* is unchanged in the Rhl-negative background compared to the wildtype. (D) The expression of the QS-regulator gene *rhlR* is reduced in the Las-negative background. Expression levels were standardised for bacterial load. Grey shaded areas depict background fluorescence (mean +/- standard deviation) of worms exposed to the non-fluorescent, non-pathogenic *E.coli* OP50. N = number of worms form four independent experiments. * = p < 0.05; ** = p < 0.01, *** = p < 0.001, n.s. = not statistically significant.

### Virulence-factor-negative mutants show trait-specific deficiencies in host colonization

To examine whether the ability to produce shared virulence factors is important for initial bacterial uptake and host colonization, we exposed *C. elegans* to five isogenic mutants of PAO1-*mCherry*, either impaired in the production of pyoverdine (Δ*pvdD*), pyochelin (Δ*pchEF*), both siderophores (Δ*pvdDΔpchEF*), the QS receptor LasR (Δ*lasR*), or the QS receptor RhlR (Δ*rhlR*). After the feeding phase, the bacterial load of the wildtype and all three siderophores mutants were equally abundant inside hosts, whereas bacterial load was significantly reduced for the two QS-mutants compared to the wildtype (Figure 4A; ANOVA, significant variation among strains F_5,736_ = 10.50, p < 0.001; post-hoc Tukey test for multiple comparisons: p > 0.05 for all siderophore mutants, p = 0.021 for PAO1Δ*lasR*, p < 0.001 for PAO1Δ*rhlR*).

**Figure 4.**
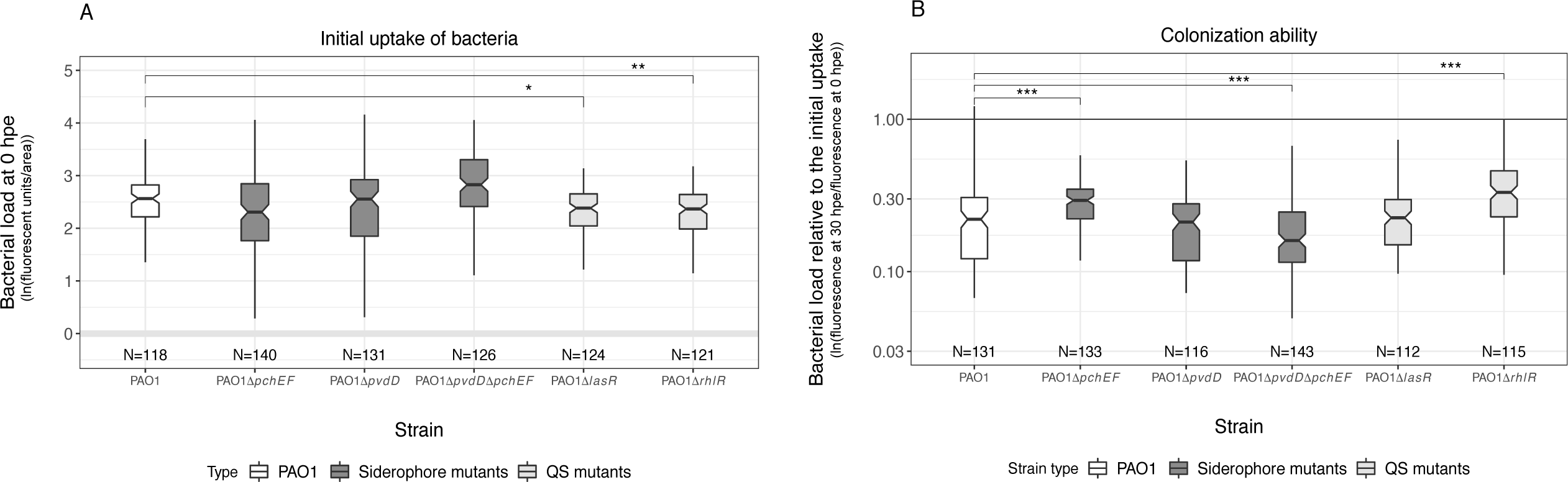
Virulence factor production affects bacterial uptake and host colonization ability. (A) Bacterial load inside *C. elegans* guts measured immediately after the recovery of worms from the exposure plates (0 hours post exposure; hpe). Comparisons across isogenic PAO1 mutant strains, each deficient for the production of one or two virulence factors, reveal that the two quorum-sensing mutants PAO1Δ*lasR* and PAO1Δ*rhlR* reached lower bacterial densities than the wildtype. (B) Comparison of the relative colonization success of strains (ratio of bacterial loads at 0 hpe versus 30 hpe) revealedthat the siderophore-negative strain PAO1Δ*pvdD*Δ*pchEF* showed significantly reduced ability to remain in the host compared to the wildtype. In contrast, the colonisation success of PAO1Δ*pchEF* and PAO1Δ*rhlR* was increased relative to the wildtype. Grey shaded areas depict background fluorescence (mean +/- standard deviation) of worms exposed to the non-fluorescent, non-pathogenic *E.coli* OP50. N = number of worms form four independent experiments. * = p < 0.05; ** = p < 0.01, *** = p < 0.001.

As previously described for PAO1 colonization (Figure 1D), we observed that the bacterial load of all strains declined at 6 hpe (Supplementary Figure S4) and 30 hpe (Figure 4B) following worm removal from the exposure plates. This decline was significantly more pronounced for the double-siderophore knockout PAO1Δ*pvdD*Δ*pchEF* than for the wildtype (Figure 4B; ANOVA, post-hoc Tukey test p < 0.001). In contrast, mutants deficient in pyochelin (PAO1Δ*pchEF*) and RhlR (PAO1Δ*rhlR*) production showed a significantly higher ability to remain in the host than the wildtype (Figure 4B; ANOVA, post-hoc Tukey test p <0.001 for both strains). Taken together, our findings suggest that the two siderophores can complement each other, and that only the siderophore double mutant and the LasR-deficient strain have an overall disadvantage in colonizing worms.

### Mixed communities are formed inside hosts, but exploitation of social traits is constrained

Given our findings on colonization deficiencies, we reasoned that the siderophore-double mutant (PAO1Δ*pvdD*Δ*pchEF*) and the Las-deficient mutant (PAO1Δ*lasR*) could act as cheats and benefit from the exploitation of virulence factors produced by the wildtype in mixed infections. To test this hypothesis, we first competed the PAO1-*mcherry* strain against the untagged wildtype in the host, and found that the mCherry tag had a small negative effect on PAO1 fitness (Figure 5A; one sample t-test, t_24_ = −4.12, p < 0.001). We then competed PAO1-*mCherry* against the two putative cheats and found that neither of them could gain a significant fitness advantage over the wildtype, but also did not lose out (Figure 5A; ANOVA, F_2,70_ = 0.517, p = 0.598). These results indicate that virulence-factor-negative mutants, initially compromised in host colonization, can indeed benefit from the presence of the wildtype producer, but not to an extent that would allow them to increase in frequency and displace producers.

**Figure 5.**
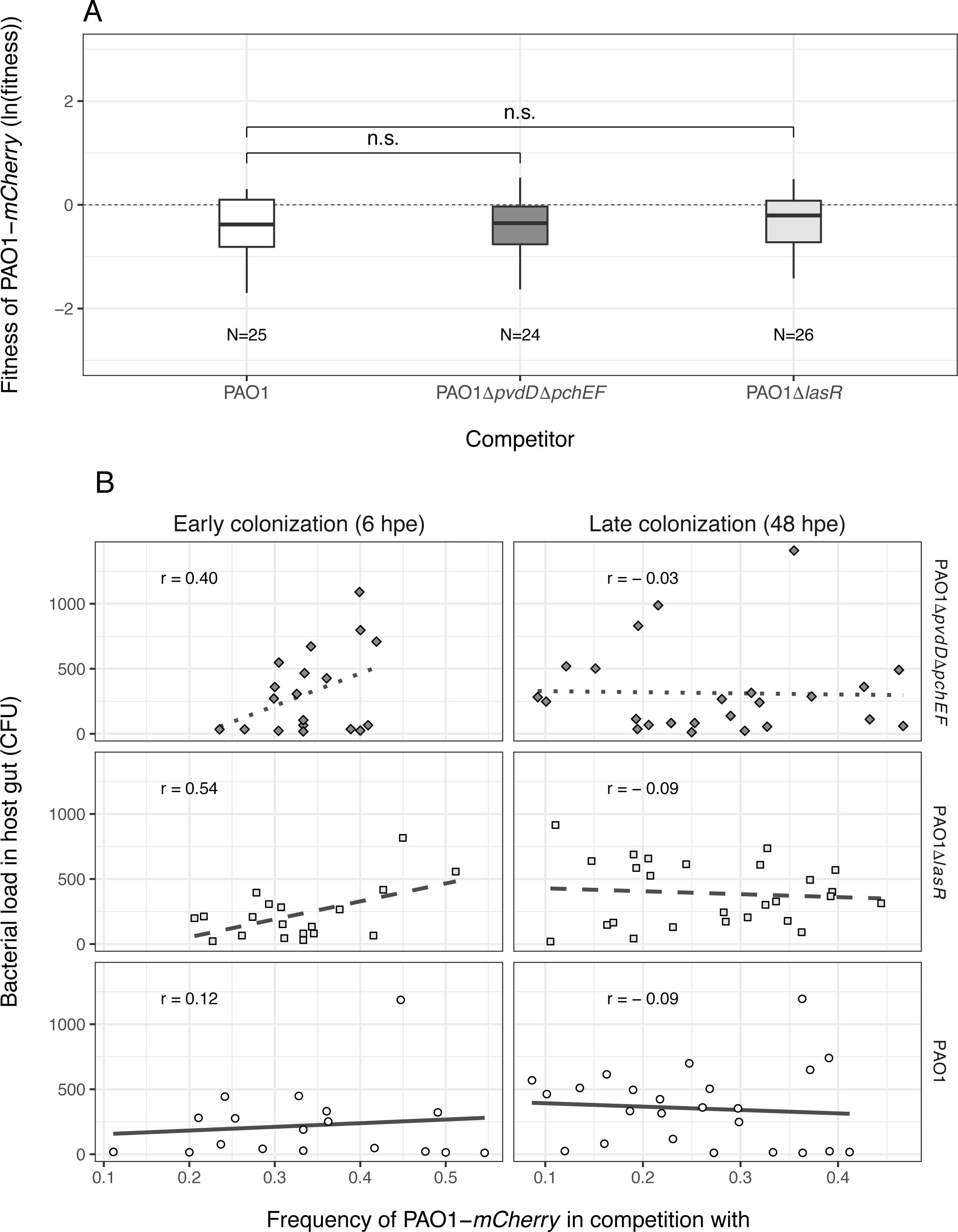
Mixed infections reveal social strain dynamics but no successful cheating. (A) Relative fitness of the wildtype PAO1-*mCherry* after 42 hours of competition inside the *C. elegans* gut against an untagged PAO1 control strain; the siderophore-negative strain PAO1Δ*pvdD*Δ*pchEF*; and the Las-negative strain PAO1Δ*lasR*. The control competition revealed a mild but significant negative effect of the mCherry tag on wildtype fitness. When accounting for these mCherry costs, we found that the putative cheat strains PAO1 Δ*lasR* and PAO1 Δ*pvdD*Δ*pchEF* performed equally well compared to the wildtype, but could not outcompete it. This suggests that virulence factor deficient strains benefit from the presence of non-producers but cannot successfully cheat on them. (B) At 6 hpe, wildtype frequency in mixed infections correlated positively with total bacterial load inside hosts in competition with PAO1Δ*lasR* (squares and dashed lines) and PAO1Δ*pvdD*Δ*pchEF* (diamonds and dotted lines) but not in the control competition (circles and solid lines). These correlations disappeared at 48 hpe. Each data point represents an individual worm. Data shown in A+B stem from the same three independent experiments.

Since our mono-infection experiments showed that the wildtype can maintain higher bacterial loads in the worms compared to the two mutants (Figure 4B), we hypothesized that worms, which have initially taken up higher frequencies of the wildtype relative to the mutant should carry increased bacterial loads in the gut. We found this prediction to hold true at 6 hpe in mixed infections with the two non-producers, but not in the control mixed infections with the untagged wildtype (Figure 5B, 6hpe; Pearson correlation coefficient, for mixed infection with PAO1Δ*lasR*: r = 0.54, t_17_ = 2.67, p = 0.016; with PAO1Δ*pvdD*Δ*pchEF*: r = 0.40, t_17_ = 1.77, p = 0.031; with control PAO1: r = 0.12, t_17_ = 0.47, p = 0.639). These correlations disappeared at the later colonization stage (Figure 5B, 48 hpe; Pearson correlation coefficient r < 0, p > 0.05 for all strains). The loss of these correlations indicates that rare producers experienced a selective advantage during competition and increased in relative frequency, while common producers might have lost and decreased in frequency.

### Strain co-localization is generally high within the host, but varies substantially across individuals

*In-vitro* studies have shown that spatial proximity of cells is crucial for efficient compound sharing [59, 60]. We thus assessed the co-localization of strains in mixed infections inside the gut (Figure 6). We found that all worms were colonized by both strains, and that the level of co-localization ***ρ*** (from tail to head) was generally high, although it varied substantially across individuals (Figure 6A-B). Similar co-localization patterns emerged for all three strain combinations tested, highlighting that the type of competitor did not influence the degree of strain co-localization in the host gut (Figure 6C; ANOVA, F_2,102_ = 2.17, p = 0.119). While our measure of co-localization has some limitations as it does not reveal physical proximity at the single cell level, and is based on a 2D projection of a 3D organ, it clearly suggests that competing cells are close to one another, and that social interactions could occur between them.

**Figure 6.**
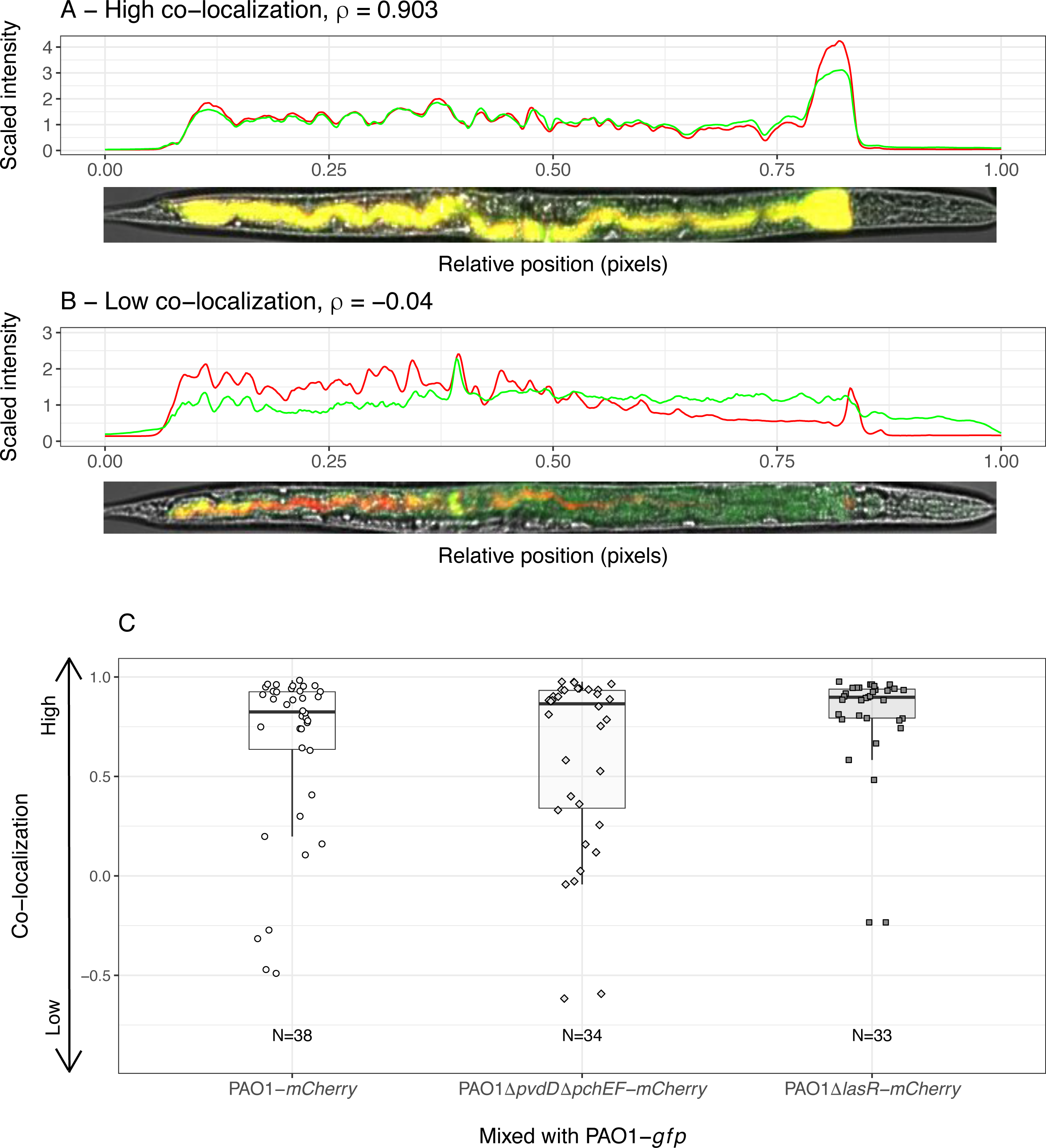
Spatial structure of mixed infections in the nematode gut. (A, B) Illustrative examples of *C. elegans* individuals infected with a mixture of GFP- and mCherry-labelled strains. Each worm was computationally straightened and fluorescence intensity values were extracted for each pixel from tail (X=0) to head (X=1). We then calculated the Spearman correlation coefficient *ρ* between the intensity values in the two fluorescence channels across pixels, as our estimate of strain colocalization. Examples show worms with high (A) and low (B) degrees of colocalization. (C) Patterns of colocalization levels varied substantially between individuals, but did not differ across strain combinations (p = 0.119: wildtype PAO1-*mCherry* versus: (i) wildtype PAO1-*gfp* (circles), (ii) PAO1Δ*pvdD*Δ*pchEF*-*mCherry* (diamonds) or (iii) PAO1Δ*lasR*-*mCherry* (squares). Each data point represents an individual worm. Data stems from 3 independent experiments, with 12 replicates each.

## Discussion

We developed a live imaging system that allows us to track host colonization by pathogenic bacteria (*P. aeruginosa*) and their expression of virulence factors inside hosts (*C. elegans*). We used this system to focus on the role of secreted virulence factors, which can be shared as public goods between bacterial cells, and examined competitive dynamics between virulence factor producing and non-producing strains in the host. We found that siderophores (pyoverdine and pyochelin) and the Las and Rhl QS-systems (i) are expressed inside the host; (ii) affect the ability to colonize and reside within the nematodes; (iii) allow non-producers to benefit from virulence factors secreted by producers in mixed infections; but (iv) do not allow non-producers to cheat and outcompete producers. Our results have implications for both the understanding of bacterial social interactions within hosts, and therapeutic approaches that aim to manipulate social dynamics between strains for infection control.

Numerous *in*-*vitro* studies have shown that bacterial cooperation can be exploited by cheating mutants that no longer express the social trait, but benefit from the cooperative acts performed by others [3, 11, 12, 55, 61–65]. These findings contrast with our observations that the spread of non-producers was constrained within infections. There are multiple ways to explain this constraint. First, increased spatial structure can limit the diffusion of secreted metabolites and lead to the physical separation of cooperators and cheats [63]. Both effects result in metabolites being shared more locally among cooperative individuals. The physical separation of strains seemed to explain the results of Zhou *et al* [20], where QS-mutants of *Bacillus thuringiensis* infecting caterpillars could not exploit metabolites from producers. Conversely, physical separation seemed low in our study system (Figure 6), and therefore unlikely explains why cheats could not spread. While we solely focussed on proximity patterns inside hosts, it is important to note that processes at the meta-population level such as bottlenecking [54, 66], can reduce the probability of different strains ending up in the same host, and thus further compromise cheat success.

Second, negative frequency-dependent selection could explain why the spread of virulence factor negative mutants is constrained [55]. This scenario predicts that cheats only experience a selective advantage when rare, but not when common. The reasoning is that non-producers can only efficiently exploit public goods when surrounded by many producers. Our competition experiments indeed provide indirect evidence for negative frequency-dependent selection in the nematode gut (Figure 5B). Specifically, we observed that bacterial load was reduced when producers occurred at low frequency early during infection (6 hpe), a result confirming that non-producers are worse host colonizers than producers. These correlations disappeared during the competition period (48 hpe), indicating that rare producers might have experienced a selective advantage and increased in relative frequency, while common producers lost and decreased in frequency.

Third, the relatively low bacterial density in the gut could further compromise the ability of non-producers to cheat (Figure 1D, 5B). Low cell density restricts the sharing and exploitation of secreted compounds [67, 68]. Mechanisms responsible for the low bacterial density in the gut (Figure 1D, 5B) could include the peristaltic activity of the gut, expelling a part of the pathogen population and the host immune system, killing a fraction of the bacteria [69].

Fourth, our analysis reveals that, although siderophores and QS-systems play a role in host colonization, they are not essential (Figure 4). Moreover, the expression of pyoverdine and QS-systems declined over time (Figure 2). These two observations indicate that the benefits of cheating might be fairly low, and that the costs of virulence factor production are reduced at later stages of the infection. Thus, bacteria might switch from production to recycling of already secreted public goods [70, 71], an effect that can hamper the spread of cheats.

Finally, we show that the regulatory linkage between traits is an important factor to consider when predicting the putative advantage of non-producers [72, 73]. For instance, *P. aeruginosa* pyoverdine-negative-mutants upregulated pyochelin production to compensate for the lack of their primary siderophore (Figure 3). Thus, if pyoverdine-negative mutants evolve *de novo*, their spread as cheats could be hampered because they invest in pyochelin as an alternative siderophore [74]. For QS, meanwhile, we observed that the absence of a functional Las-system resulted in the concomitant collapse of the Rhl-system. Although *lasR* mutants could be potent cheats, as they are deficient for multiple social traits, their spread might be hampered because QS-systems also regulate non-social traits, important for individual fitness [75].

When relating our insights to previous studies, it turns out that earlier work produced mixed results with regard to the question whether siderophore- and QS-deficient mutants can spread within infections. While Harrison *et al*. ([15, 21]; pyoverdine, *P. aeruginosa* in *Galleria mellonella* and *ex-vivo* infection models) and Zhou *et al*. ([20]; QS, *B. thuringiensis* in *Plutella xylostella*) showed that the spread of non-producers is constrained, Rumbaugh *et al*. ([16, 17]; QS; *P. aeruginosa* in mice), Pollitt *et al*. ([19]; QS, *Staphylococcus aureus* in *G. mellonella*) and Diard *et al*. ([18], T3SS-driven inflammation, *Salmonella typhimorium* in mice) demonstrated cases where non-producers spread to high frequencies in host populations. While the reported results were based on strain frequency counts before and after competition, we here show that information on social trait expression, temporal infection dynamics and physical interactions among strains within hosts are essential to understand whether social traits are important and exploitable in a given system. We thus posit that more such detailed approaches are required to understand the importance of bacterial social interactions across host systems and infection contexts and explain differences between them.

A deeper understanding of bacterial social interactions inside hosts is particularly relevant for novel therapeutic approaches that seek to take advantage of cooperator-cheat dynamics inside hosts to control infections. For instance, it was proposed that strains deficient for virulence factors could be introduced into established infections [24]. These strains are expected to spread because of cheating, thereby reducing the overall virulence factor availability in the population and the damage to the host. Our results reveal that virulence-factor-negative strains, although eventually gaining a benefit from producer strains, are unable to spread in populations. Another therapeutic approach involves the specific targeting of secreted virulence factors [25, 27]. This approach is thought to reduce damage to the host and to compromise resistance evolution [30]. Resistant mutants, resuming virulence factor production, are not expected to spread because they would act as cooperators, sharing the benefit of secreted goods with susceptible strains [76–78]. Our results yet indicate that such resistant mutants could get local benefits and thus increase to a certain frequency in the population [31]. These confrontations show that the identification of key parameters driving social interactions across hosts and infection types is of utmost importance to predict the success of ‘cheat therapies’ and anti-virulence strategies targeting secreted public goods.

## Acknowledgments

We thank two anonymous reviewers for constructive comments and the Center of Microscopy and Image Analysis (University of Zürich) for support with image acquisition and advice on image analysis.

## Supplementary Information

Supplementary information is available at the journal’s website.

