## Supplementary figures and images for "*In vivo* microscopy reveals the impact of *Pseudomonas aeruginosa* social interactions on host colonization"

### Supplementary Figure S1

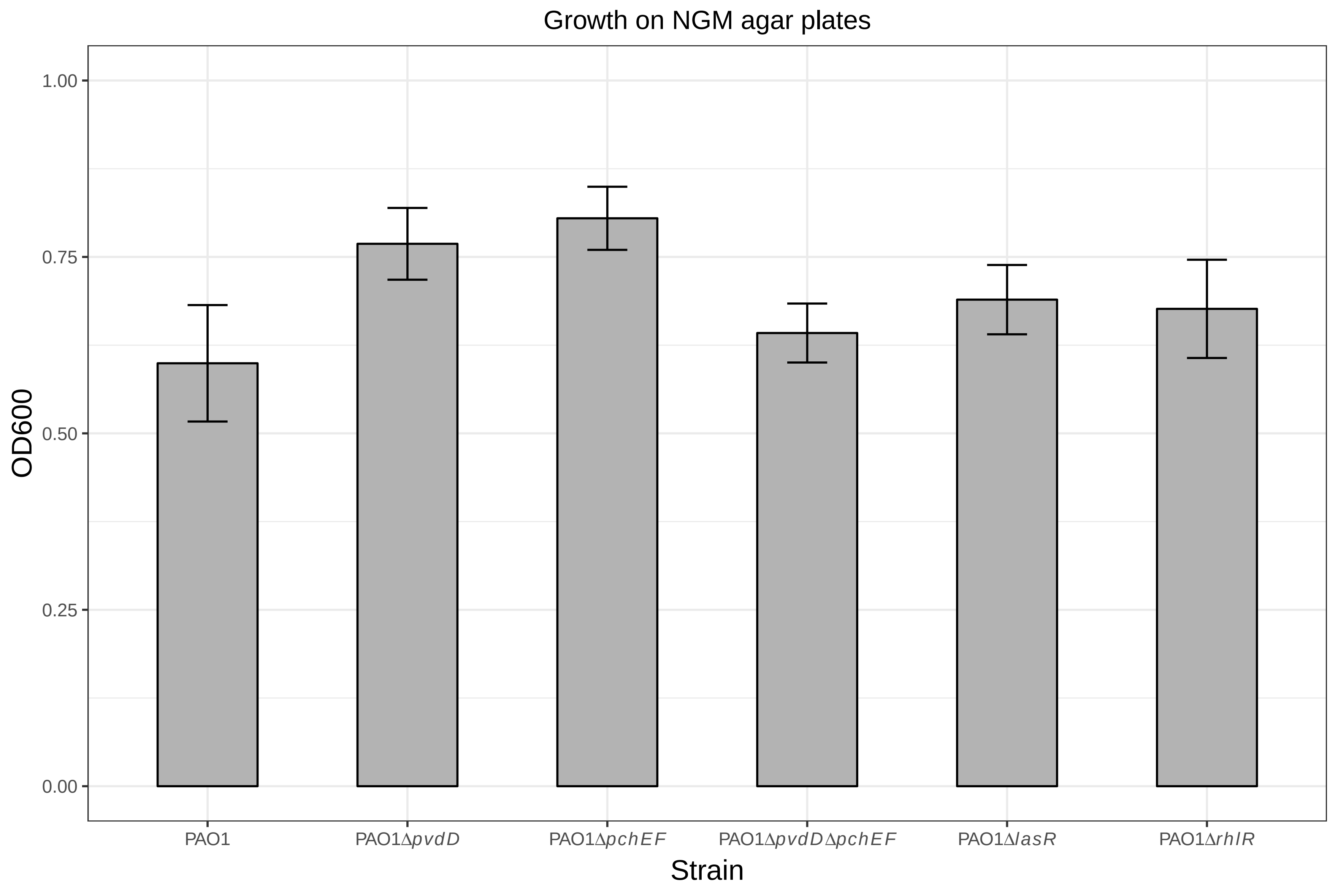

### Supplementary Figure S2

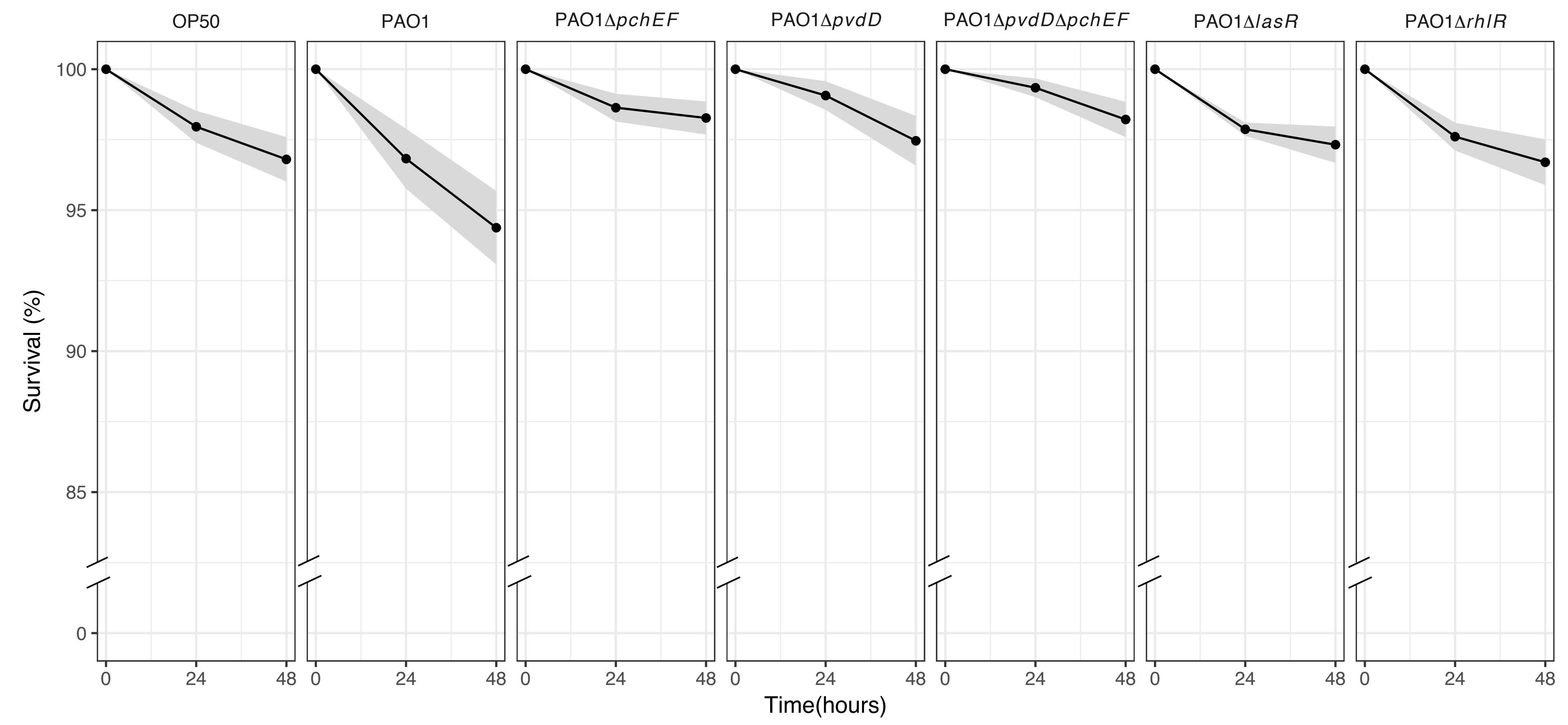

### Supplementary Figure S3

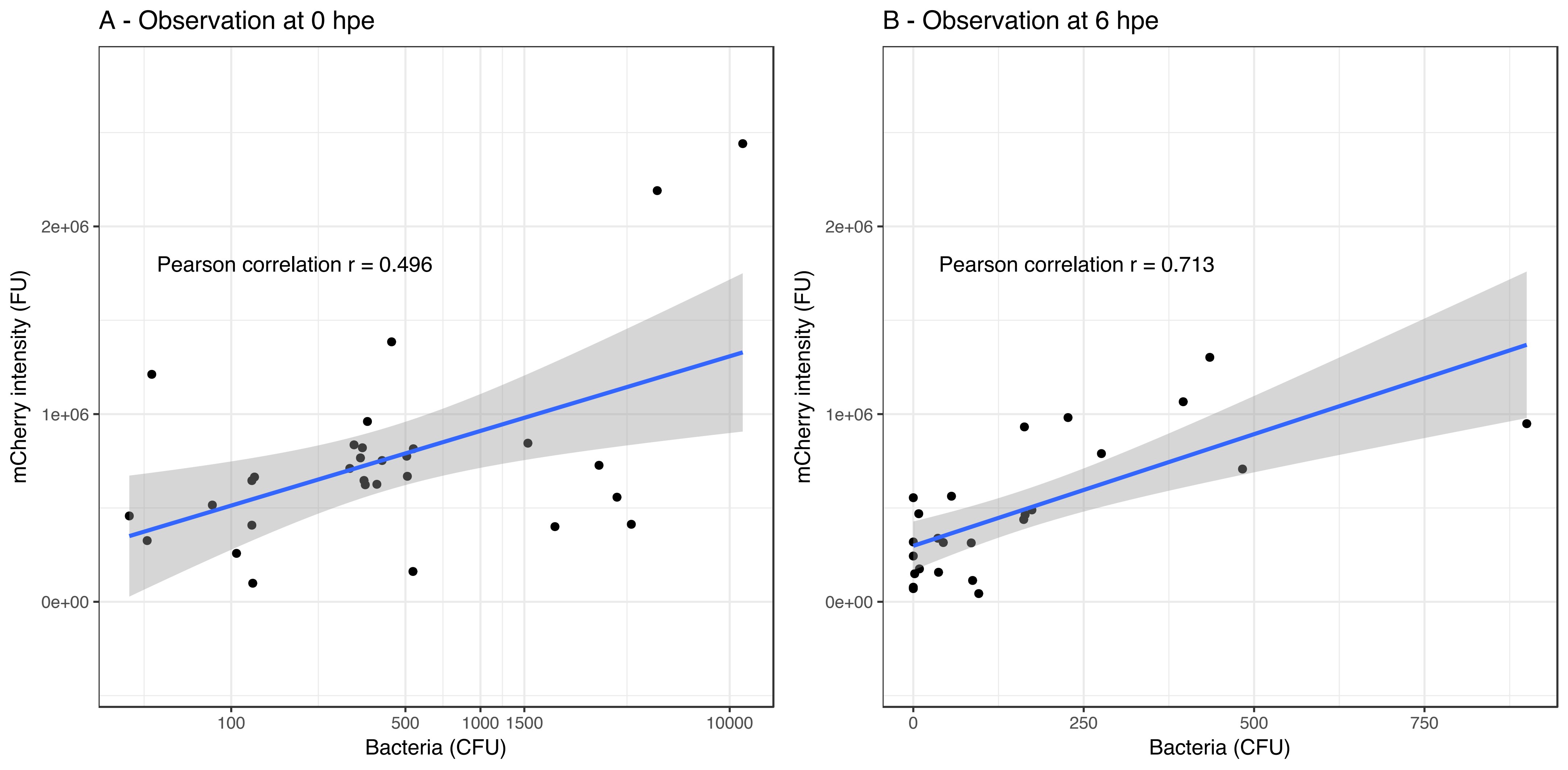

### Supplementary Figure S4

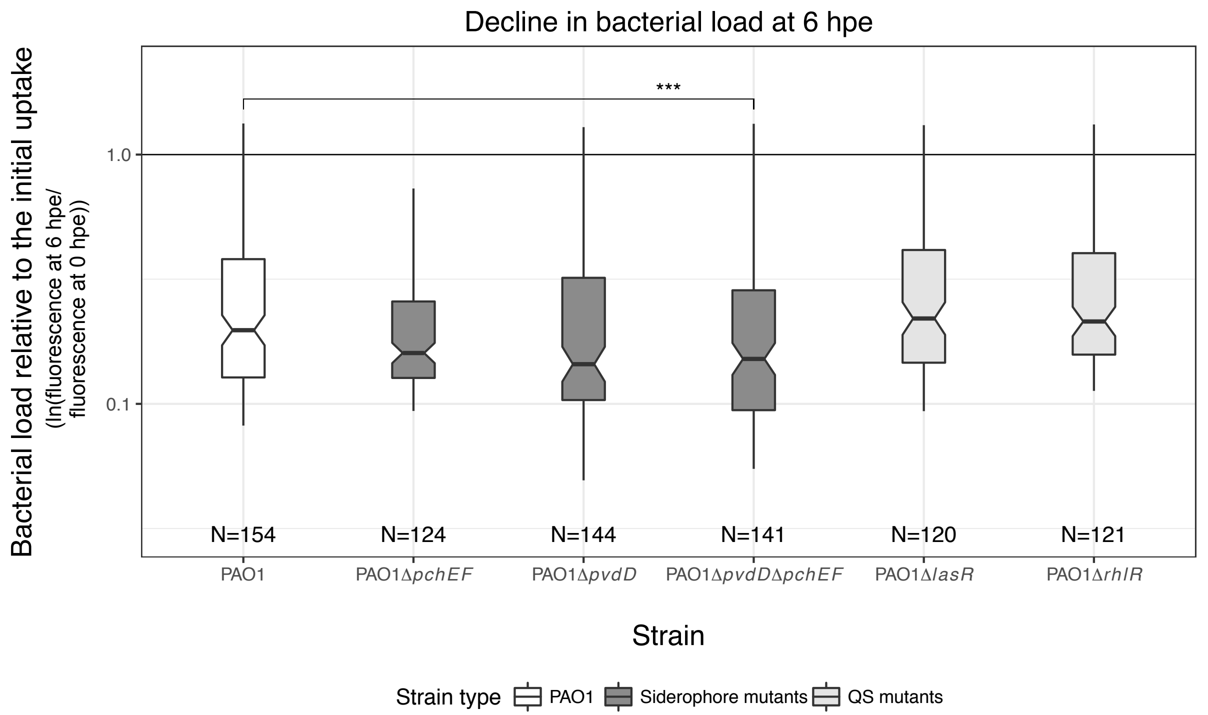

### Supplementary Figure S5

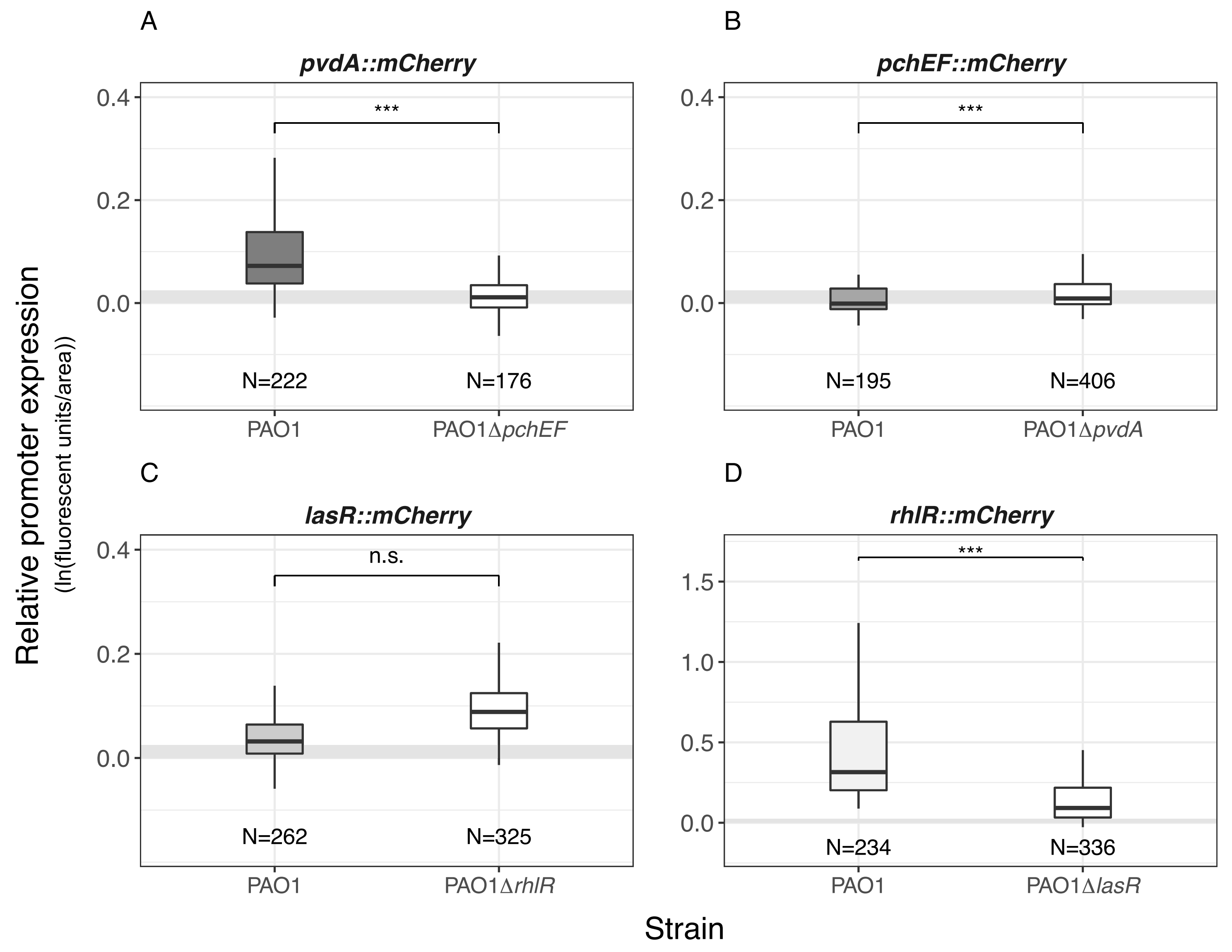
