## Supplementary Figure Captions for "*In vivo* microscopy reveals the impact of *Pseudomonas aeruginosa* social interactions on host colonization"

**Supplementary Figures captions**

**Supplementary Figure S1. Growth of *P. aeruginosa* WT and mutant strains for social traits on NGM plates.** To assess the growth ability of PAO1 and the mutant strains on NGM plates, after 24 hours of incubation at 25°C, we collected the bacterial lawn in sterile NaCl solution and the OD600 was measured as proxy for cell growth. We found no statistically significant difference between strains (linear model F_5,60_ = 2.27, p = 0.0588). Data is shown as mean across four independent experiments, with three replicates (i.e. plates) per strain for each experiment. Error bars denote the standard error of the mean.

**Supplementary Figure S2. Survival assay of worms infected with various *P. aeruginosa* strains.** After washing worms off the exposure plates, we estimated host survival over 48 hours. For this purpose, we observed 50 to 90 worms and checked for viability every 24 hours by prodding them with a platinum wire three times. Worms were considered dead if they no longer moved upon stimulation with the wire . We found no significant difference in the survival rate between any of the strains compared to the food strain *E.coli* OP50 (linear model, F_6,210_ = 0.60, p = 0.7296). However, we found a small but significant difference in the survival of worms colonized by the three siderophore-mutants, which was higher compared to the survival of worms exposed to PAO1 (Tukey's range test, p = 0.0213 for PAO1Δ*pchEF*, p = 0.0054 for PAO1Δ*pvdD* and p = 0.0062 for PAO1Δ*pvdD*Δ*pchEF*). Data points depict average survival across three independent experiments. In each experiment, we had three replicates for each strain. Gray areas represent the standard error of the mean.

**Supplementary Figure S3. Fluorescence intensity significantly correlates with live bacteria inside the host gut.** To assess the relationship between fluorescence signal and bacterial load inside *C. elegans*, colonized nematodes were observed using fluorescence microscopy and disrupted to extract live bacteria from the gut. Fluorescence intensity significantly correlated with the number of bacteria present in the host gut. (A) The correlation was moderate when the worms were observed immediately after exposure (0 hours post exposure; hpe) (Pearson correlation coefficient r = 0.496; test for association between paired samples t_28_ = 3.02, p = 0.0053). (B) At 6 hpe, fluorescence intensity correlated more strongly with bacterial load in the host gut (Pearson correlation coefficient r = 0.713; test for association between paired samples t_23_ = 4.88, p < 0.0001). In total, 65 worms were observed in two independent experiments. Fluorescence intensity values were blank corrected, using worms infected with the untagged strain PAO1 as non-fluorescent controls.

**Supplementary Figure S4. Bacterial load declines at 6 hours post exposure (hpe).** We kept infected nematodes in sterile buffer and determined the fluorescence intensity in the host gut, reflecting the number of bacteria that managed to colonize the worms at 6 hpe. When scaled to the bacterial load at 0 hpe, we found that all strains showed a significant reduction in bacterial load. Moreover, the siderophore-negative strain PAO1Δ*pvdD*Δ*pchEF* showed significantly reduced ability to remain in the host compared to PAO1 (ANOVA with post-hoc Tukey test, t_888_ = -2.25, p = 0.0025). For each individual strain, relative fluorescence is expressed as fluorescence intensity at 6 hpe scaled for the intensity at 0 hpe.

**Supplementary Figure S5. Interaction between social traits inside the host at 30 hours post exposure.** We compared the expression of promoter fusions inserted either in the wildtype PAO1 or in mutant strains, which lack either the second siderophore or the second QS-regulator (white boxplots), at 30 hpe. Although values are generally lower, we observed the same trend as in Figure 3. Values are corrected for cell density. N = number of worms tested. Error bars represent standard errors of the mean. Grey shaded areas indicate the non-fluorescent background (mean +/- standard deviation). *** = p < 0.001 and n.s. = not statistically significant, based on Welch’s 2-sample t-test between PAO1 and the respective mutant strain.
