## Supplementary Methods for "*In vivo* microscopy reveals the impact of *Pseudomonas aeruginosa* social interactions on host colonization"

**Promoter-mCherry fusions construction**

Bacterial strains used in this study are listed in Supplementary Table S1. To construct reporter gene fusions, chromosomal DNA of PAO1 was extracted using the *GenElute Bacterial Genomic DNA Kit* (Sigma-Aldrich, Switzerland) following the instructions of the manufacturer. Promoter regions of genes of interest (*pvdA*, *pchEF*, *lasR*, *rhlR*) were amplified using *Phusion High-Fidelity DNA Polymerase* (New England BioLabs, Switzerland). Primers and conditions used to amplify the promoter regions are described in Supplementary Table S2. PCR products were purified using *the GeneElute PCR Clean-Up Kit* (Sigma-Aldrich) and digested using the appropriate restriction enzymes (Thermo Fischer Scientific, Switzerland). DNA fragments were purified using *Select-a-Size DNA Clean & Concentrator* (Zymo Research, California, United States).

PCR products were then inserted into an empty pUC18-miniTn7-Gm vector [1], using a T4 Ligase enzyme (Thermo Fischer Scientific) and transformed in *Escherichia coli* SM10λpir by the standard CaCl_2_ procedure [2]. Transformants were selected on LB + Ampicillin (100 μg/mL) and confirmed by colony PCR using the primer “Tn7L_rev”, annealing on the vector backbone and each “Promoter_down” specific primer (Supplementary Table S2). Colony PCR was performed using the *OneTaq Hot Start DNA polymerase* (New England BioLabs). All cloning steps were verified by DNA Sanger sequencing, which was performed by Microsynth (Switzerland). Plasmids containing promoter-mCherry fusions were then extracted using the *QIAprep Spin Miniprep Kit* (QIAGEN, Switzerland) and are listed in Supplementary Table S3.

Transformation of *P. aeruginosa* was performed by electroporation following the protocol described in [1]. The mini-Tn7 system ensures single-copy chromosomal insertion of the fluorescent reporter in the *att*Tn7 site of *P. aeruginosa*. Transformants were selected on LB + Gentamycin (30 μg/mL) and confirmed by colony PCR, using the primer pair “PglmS_up” and “Tn7L”.
