## Supplementary Table S1 for "*In vivo* microscopy reveals the impact of *Pseudomonas aeruginosa* social interactions on host colonization"

**Supplementary Table S1. List of strains used in this study**

| **Strain name** | **Description or genotype** | **Source or reference** |
| --- | --- | --- |
| ***E. coli*** |  |  |
| SM10λpir | *thi thr leu tonA lacY supE recA*::RP4-2-Tc::Mu λ*pir*R6K Km^r^ | [1] |
| OP50 | Food source for *C. elegans* | Caenorhabditis Genetic Center |
| ***P. aeruginosa*** |  |  |
| PAO1 | Wild type strain | This laboratory |
| PAO1-*mCherry* | PAO1 with constitutive mCherry expression (*att*Tn7::*Ptac-mCherry)* | This laboratory |
| PAO1-gfp | PAO1 with constitutive GFP expression (*att*Tn7::*Ptac-GFP)* | This laboratory |
| PAO1*ΔpvdD-mCherry* | Deletion mutant, deficient for pyoverdine production and with constitutive mCherry expression (*att*Tn7::*Ptac-mCherry)* | This laboratory |
| PAO1*ΔpchEF-mCherry* | Deletion mutant, deficient for pyochelin production and with constitutive mCherry expression (*att*Tn7::*Ptac-mCherry)* | This laboratory |
| PAO1*ΔpvdDΔpchEF* | Deletion mutant, deficient for pyoverdine and pyochelin production | This laboratory |
| PAO1*ΔpvdDΔpchEF-mCherry* | Deletion mutant, deficient for pyoverdine and pyochelin production, with constitutive mCherry expression (*att*Tn7::*Ptac-mCherry)* | This laboratory |
| PAO1*ΔlasR* | Deletion mutant, deficient for the production of the transcriptional regulator LasR | S. Diggle strain collection, Georgia Tech |
| PAO1*ΔlasR-mCherry* | Deletion mutant, deficient for the production of the transcriptional regulator LasR and with constitutive mCherry expression (*att*Tn7::*Ptac-mCherry* from pUC18-miniTn7-Gm-*mCherry*) | This study |
| PAO1*ΔrhlR* | Deletion mutant, deficient for the production of the transcriptional regulator RhlR | S.Diggle strain collection, Georgia Tech |
| PAO1*ΔrhlR-mCherry* | Deletion mutant, deficient for the production of the transcriptional regulator RhlR and with constitutive mCherry expression (*att*Tn7::*Ptac-mCherry* from pUC18-miniTn7-Gm-*mCherry*) | This study |
| PAO1-*pvdA-mCherry* | PAO1 with the transcriptional fusion *pvdA::mCherry* from pCR01 | This study |
| PAO1-*pchEF-mCherry* | PAO1 with the transcriptional fusion *pchE::mCherry* from pCR02 | This study |
| PAO1-*lasR-mCherry* | PAO1 with the transcriptional fusion *lasR::mCherry* from pCR03 | This study |
| PAO1-*rhlR-mCherry* | PAO1 with the transcriptional fusion *rhlR::mCherry* from pCR04 | This study |
| PAO1*ΔpvdD, pchEF-mCherry* | Deletion mutant, deficient for pyoverdine production, with the transcriptional fusion *pchE::mCherry* from pCR02 | This study |
| PAO1*ΔpchEF, pvdA-mCherry* | Deletion mutant, deficient for pyochelin production, with the transcriptional fusion *pvdA::mCherry* from pCR01 | This study |
| PAO1*ΔrhlR, lasR-mCherry* | Deletion mutant, deficient for the production of the transcriptional regulator RhlR, with the transcriptional fusion *lasR::mCherry* from pCR03 | This study |
| PAO1*ΔlasR, rhlR-mCherry* | Deletion mutant, deficient for the production of the transcriptional regulator LasR, with the transcriptional fusion *rhlR::mCherry* from pCR04 | This study |
