## Supplementary Table S2 for "*In vivo* microscopy reveals the impact of *Pseudomonas aeruginosa* social interactions on host colonization"

**Supplementary Table S2. List of primers used in this study**

| **Primer name** | **Sequence (5’-3’)** | **Application** | **Template** | **T_a_^1^** |
| --- | --- | --- | --- | --- |
| pvdA_down_SacI | CGG CAT CAG AGC AGA TTG TA | Plasmid pCR01 | pEX-pvdA-*mCherry* | 60°C |
| pvdA_up_HindIII | GGA TCC AAG CGA GCA AAA G | Plasmid pCR01 | pEX-pvdA-*mCherry* | 60°C |
| pchEF_down_BamHI | CGA GGG ATC CTC ACT GCT CGG TCA GCC AGT C | Plasmid pCR02 | PAO1 gDNA | 54°C |
| pchE_up_HindIII | GAT CAA GCT TCA AGC GCT ACG GCA TCT C | Plasmid pCR02 | PAO1 gDNA | 54°C |
| lasR_down_BamHI | CGT TCG GAT CCT TAG GCG CTC CAC TCC AAT TTT C | Plasmid pCR03 | PAO1 gDNA | 65°C |
| lasR_up_HindIII | ATG ACA AGC TTT GGA AAA GTG GCT ATG TCG C | Plasmid pCR03 | PAO1 gDNA | 65°C |
| rhlR_down_BamHI | TTG CTG GAT CCT CAC TGC ATC TGG TAT CGC TCC | Plasmid pCR04 | PAO1 gDNA | 63°C |
| rhlR_up_HindIII | TAA CGA AAG CTT CCT GCA GGG CGA CTT CTA C | Plasmid pCR04 | PAO1 gDNA | 63°C |
| Tn7L_rev | GGG TGT AGC GTC GTA AGC TAA T | Colony PCR *E. coli* | *E. coli* colonies | 53-54°C^2^ |
| PglmS_up | CTG TGC GAC TGC TGG AGC TGA | Colony PCR in PAO1 | PAO1 colonies | 54°C |
| Tn7L | ATT AGC TTA CGA CGC TAC ACC C | Colony PCR in PAO1 | PAO1 colonies | 54°C |

^1^ Annealing temperature used to amplify promoter regions

^2^ To confirm the insertion of the promoter region into the miniTn7 vector, colony PCR was performed using the Tn7L_rev primer, together with the corresponding “down” promoter-specific primer. Annealing temperatures were: 53°C for *pvdA*; 54°C for *pchEF*, *lasR*; *rhlR*.
