## Supplementary Table S3 for "*In vivo* microscopy reveals the impact of *Pseudomonas aeruginosa* social interactions on host colonization"

**Supplementary Table S3. List of plasmids used in this study**

| **Plasmid name** | **Description** | **Source or reference** |
| --- | --- | --- |
| pUX-BF13 | Helper plasmid, providing the Tn7 transposase proteins | [1] |
| pUC18-miniTn7-Gm | Gm^r^ on mini-Tn7; for gene insertion in Gm^s^ bacteria | [2] |
| pUC18-miniTn7-Gm-*mCherry* | Gm^r^ on mini-Tn7; for *Ptac::mCherry* tagging in Gm^s^ bacteria in the *att*Tn7 site | [2] |
| pEX-pvdA-*mCherry* | Commercial plasmid with *pvdA::mCherry* between SacI/HindIII sites | [3] |
| pRC01 | pUC18-mini-Tn7-Gm with *pvdA::mCherry* from pEX-pvdA-*mCherry* (SacI/HindIII) | This study |
| pRC02 | pCR01, with *pvdA* promoter replaced by *pchEF* promoter (BamHI/HindIII) | This study |
| pRC03 | pCR01 with *pvdA* promoter replaced by *lasR* promoter (BamHI/HindIII) | This study |
| pRC04 | pCR01 with *pvdA* promoter replaced by *rhlR* promoter (BamHI/HindIII) | This study |
